# *Legionella* relative abundance in shower hose biofilms is associated with specific microbiome members

**DOI:** 10.1101/2023.05.04.539404

**Authors:** Alessio Cavallaro, William J. Rhoads, Émile Sylvestre, Thierry Marti, Jean-Claude Walser, Frederik Hammes

## Abstract

*Legionella* are natural inhabitants of building plumbing biofilms, where interactions with other microorganisms influence their survival, proliferation, and death. Here, we investigated the associations of *Legionella* with prokaryotic and eukaryotic microbiomes in biofilm samples extracted from 85 shower hoses of a multiunit residential building. *Legionella* spp. relative abundance in the biofilms ranged between 0 - 7.8%, of which only 0 - 0.46% was *L. pneumophila*. Our data suggest that some microbiome members were associated with high (e.g., *Chthonomonas*, *Vrihiamoeba*) or low (e.g., *Aquabacterium*, *Vannella*) *Legionella* relative abundance. The correlations of the different *Legionella* variants (30 Zero-Radius OTUs detected) showed distinct patterns, suggesting separate ecological niches occupied by different *Legionella* species. This study provides insights into the ecology of *Legionella* with respect to: 1) the colonization of a high number of real shower hoses biofilm samples; 2) the ecological meaning of associations between *Legionella* and co-occurring prokaryotic/eukaryotic organisms; 3) critical points and future directions of microbial-interaction-based-ecological-investigations.

## 1. Introduction

The genus *Legionella* comprises over 60 species, of which about half are recognised opportunistic pathogens and the etiological agents of Legionellosis. *Legionella pneumophila* was the first species identified, and is still linked with most reported Legionellosis cases (Fields *et al*., 2002). However, multiple other *Legionella* species are ecologically relevant, and sometimes associated with different clinical manifestations of the disease (Muder & Victor, 2002, Chambers *et al*., 2021). *Legionella* are ubiquitous in freshwater, and commonly detected in engineered aquatic ecosystems such as drinking water systems (Falkinham *et al*., 2015). As such, *Legionella* have been detected in all segments of building plumbing systems, including shower hoses, which often represent a favourable environment for their proliferation and transmission (Falkinham *et al*., 2015, Proctor *et al*., 2016, Gebert *et al*., 2018). In these environments, *Legionella* mainly establish in biofilms, where interactions with other organisms influence their survival and proliferation (Declerck, 2010).

Interactions with their surrounding microbiota may be either favourable or detrimental for *Legionella* (Taylor *et al*., 2009, Cavallaro *et al*., 2022). For example, *Legionella* are able to infect a variety of eukaryotes such as amoebae, excavates, and ciliates, escaping their degradation mechanisms and exploiting their intracellular environment for protection and replication (Boamah *et al*., 2017, Mondino *et al*., 2020). However, there are also protists such as *Willaertia magna* and *Paramecium multimicronucleatum* that have been documented to degrade *Legionella* under specific environmental conditions (Dey *et al*., 2009, Amaro *et al*., 2015, Boamah *et al*., 2017). Both positive and negative interactions between *Legionella* and other prokaryotes have also been described previously. For example, *Legionella* was demonstrated to grow in association with Cyanobacteria, as well as with *Flavobacterium breve (Tison et al., 1980, Berendt, 1981, Wadowsky & Yee, 1983)*. Toze and colleagues (Toze *et al*., 1990) observed that heterotrophic bacteria isolated from chlorinated drinking water were able to support the growth of some species of *Legionella* but inhibited others. Others have also observed inhibition zones surrounding heterotrophic bacteria colonies isolated from various water sources when co-cultured with *Legionella* (Guerrieri *et al*., 2008, Corre *et al*., 2018). A few recent studies investigated correlations between *Legionella* and other microbiome members identified through amplicon sequencing. Paranjape and colleagues (Paranjape *et al*., 2020, Paranjape *et al*., 2020) conducted two studies on cooling tower water and found that the genus *Pseudomonas* exhibited a strong negative correlation with *Legionella*, which, in turn, positively correlated with the genus *Brevundimonas*; the latter positive interaction has been further demonstrated in laboratory experiments with pure culture isolates. Scaturro and co-workers (Scaturro *et al*., 2022) investigated building plumbing water samples from several European cities and found positive correlations with bacteria of the genus *Nitrospira*, while observing negative associations with the genera *Pseudomonas* and *Sphingopyxis*.

Despite these correlation or inhibition observations, it remains unclear to which degree results are general or site-specific, whether amplicon sequencing provides sufficient resolution for such complex ecological interactions, and whether a mechanistic understanding of why some members support and other inhibit *Legionella* can be deduced from statistical correlations in microbiome data. Moreover, these studies also focussed exclusively on water samples, which may not accurately reflect biofilm ecology (Ji *et al*., 2017), and large sample numbers are sometimes difficult to obtain in such studies.

Here, we extracted 85 shower hose biofilms collected in a multiunit residential building with known *Legionella* contamination that shared the same incoming water, had a similar age, and were made from the same material. We used digital droplet polymerase chain reaction (ddPCR) and amplicon sequencing to identify and quantify potential associations between *Legionella*, prokaryotes, and eukaryotes in biofilm samples with similar environmental exposure.

## 2. Materials and Methods

### 2.1. Sample collection

Samples were collected from a retirement facility in Switzerland. In total, 85 shower hoses were replaced and transported to the laboratory in sealed bags. All showers-hoses were (1) from the same manufacturer, (2) of similar operational age, and (3) fed from the same building plumbing system. Unfortunately, no specific information on use habits (e.g., frequency, temperature, etc.) could be obtained due to privacy protection concerns.

### 2.2. Biofilm and DNA extraction

Biofilm extraction was performed as described previously (Proctor *et al*., 2018). Briefly, shower hoses were cut at both ends and filled with sterile glass beads (2 mm diameter) and filtered (0.2 µm) tap water. With both ends plugged, shower hoses were inverted five times in order to allow the beads to distribute internally. Then the hoses were placed in an ultrasonic bath (5 min). After sonication, the shower hoses were unplugged and the biofilm-containing-water was collected in autoclaved Schott bottles. The entire cycle was repeated 5 times, alternating from which end the water was collected. Biofilm-containing-water samples were filtered concentrated (0.2 µm polycarbonate membrane filters) and stored at −20 °C until DNA extraction. DNA extraction was carried out using an adapted version of the DNA PowerWater Kit (Qiagen), described by Vosloo and colleagues (Vosloo *et al*., 2019). Briefly, the filters were fragmented with a scalpel, inserted into 2 mL tubes and treated 1) enzymatically with the addition of Lysozyme; 2) chemically with the addition of Proteinase K; 3) mechanically using a tube shaker and the beads provided with the kit, after the addition of chloroform/isoamyl alcohol 24:1. The rest of the extraction was conducted according to the manual provided by the manufacturer of the kit.

### 2.3. ddPCR for Legionella spp. and Legionella pneumophila

*Legionella* spp. (ssrA gene) and *L. pneumophila* (mip gene) were measured using a digital droplet polymerase chain reaction (ddPCR) duplex assay with TaqMan chemistry. Gene target primers and probes were based on previously published assays (Benitez & Winchell, 2013, Rhoads *et al*., 2022). Each 25 µL reaction contained 1XPerfeCT a Multiplex ToughMix 5X (Quantabio), 0.6 µM of ssrA and 0.4 µM of mip gene forward and reverse primers, 0.15 µM of each probe, 100 nM Fluorescein (Sigma Aldrich), and 5 µL of DNA template. Primer and probe sequences, fluorophores utilized, master mix composition, and reaction conditions can be found in Supplementary Table 1. A ddPCR reaction negative control (DNAse free water) was included for each batch of master mix. A ddPCR reaction positive control (Centre National de Référence des Légionelles) was included in each thermocycling run. Droplet formation and PCR thermocycling were performed using Stilla geodes and read using a Prism6 analyser with Crystal Reader software imaging settings pre-set and optimized for PerfeCTa multiplex master mix. Droplets were analysed using Crystal Miner software. Only wells with a sufficient number of total and analysable droplets, as well as a limited number of saturated signals, were accepted according to the Crystal Miner software quality control.

### 2.4. ddPCR for 16S quantification

Total quantification of the 16S rRNA gene was carried out using a ddPCR assay with an intercalating-dye chemistry. Each 25 µL reaction contained 1X PerfeCTa Multiplex ToughMix 5X (Quantabio), 1.5X EvaGreen (Biotium), 0.8 ng/µL AlexaFluor 488, 0.1 µM of each primer and 5 µL of the DNA template. A ddPCR reaction negative control (DNAse free water) was included for each batch of master mix prepared. A ddPCR reaction positive control was included in each run. The positive control corresponded to the 16S rRNA gene sequence (V4 region) of *Escherichia coli* and was synthesized in the form of gBlock by Integrated DNA technologies (IDT). Primer sequence, master mix composition, and reaction conditions can be found in Supplementary Table 2.

### 2.5. Amplification of 16S and 18S rRNA genes for NGS library preparation and Illumina sequencing

For sequencing, the V4 region of the 16S rRNA gene and the V9 region of the 18S rRNA gene were amplified by PCR using the primers Bakt_515F - Bakt_805R (Caporaso *et al*., 2011) and EUK1391F - EUK1510R (Tsao *et al*., 2019) and the DNA was quantified by Qubit dsDNA HS Assay. Samples were diluted according to their concentration such that 0.1 to 10 ng DNA was loaded into each reaction. Two – step PCR protocol was used to prepare the sequencing library: a first amplification (target PCR) was carried out with 1X KAPA HiFi HotStart DNA polymerase (Roche), 0.3 µM of each 16S primer and 5 µL of 0.1 to 10 ng of template DNA. After amplification, the PCR products were purified with the Agencourt AMPure System (Beckman Coulter, Inc.). The second PCR (adaptor PCR) was performed with limited cycles to attach specific sequencing Nextera v2 Index adapter (Illumina). After purification, the products were quantified and checked for correct length (bp) with the High Sensitivity D1000 ScreenTape system (Agilent 2200 TapeStation). Based on their concentration, samples were divided into three libraries, which were subsequently pooled together to a final concentration of 4 nM. The Illumina MiSeq platform was used for pair-end 600 cycle (16S) and paired-end 300 cycle sequencing with 10% PhiX (internal standard) in the sequencing run. Negative controls (PCR grade water) and a positive control (For 16S library; self-made MOCK community) were incorporated. Primer sequences, master mix composition, and reaction conditions can be found in the Supplementary Table 3. Experiments and data on community composition were generated in collaboration with the Genetic Diversity Centre (GDC) of ETH Zurich.

### 2.6. Data analysis

16S and 18S rRNA sequencing data were processed on HPC Euler (ETHZ) using workflows established by the GDC, ETHZ. Detailed data processing workflows are provided in the supplementary materials. For the 16S dataset, because of the bad quality of the reverse read, only the forward read has been processed further and used for the rest of the analysis. For the 16S dataset, all R1 reads were trimmed (based on the error estimates) by 25nt, the primer region removed, and quality filtered. For the 18S dataset, all read pairs were merged, primer sites removed, and quality filtered. Ultimately, using UNOISE3 (Edgar & Flyvbjerg, 2015), sequences were denoised with error correction and chimera removal and Amplicon Sequence Variants (ASV) established. In this study, the predicted biological sequences will be referred to as Zero-Radius Operational Taxonomic Units (ZOTUs). Taxonomic assignment was performed using the Silva 16S database (v128, 16S amplicon sequencing) and PR^2^ (V4.12.0, 18S amplicon sequencing) in combination with the SINTAX classifier. Alpha-diversity (calculated with Shannon Index, which measures intra-sample diversity taking into account both richness and evenness), distance ordination and average relative abundance analysis were performed using R (version 4.2.1) and R studio (version 2022.07.2+576) using the packages “ggplot2”, “microbiomeExplorer” (through a Shiny app, (Reeder *et al*., 2021)) and the Bioconductor “phyloseq” (version 1.42.0, (McMurdie & Holmes, 2013)). SparCC correlation analysis was performed with the R package “SpiecEasi” and built as a network with Cytoscape (version 3.9.1). Random Forest analysis was performed using Microbiome Analyst (Chong *et al*., 2020). Briefly, the samples were filtered based on a low count filter (minimum count of 4 reads with 20% prevalence in samples) and a low variance filter (10% removal with the variance measured using Interquartile-range), using the Microbiome Analyst default settings. Then, the experimental variable was set (*Legionella* spp. or *L. pneumophila* relative abundance) and a Random Forest is run with 1000 trees, 7 predictors and a Randomness setting option. The contribution of each predictor to the model was calculated and reported as Mean Decrease Accuracy (MDA), which indicates how much the model loses confidence if that predictor is removed from the model itself. The Out Of Bag (OOB) error was used as a validation parameter for the accuracy of the models. In general, the OOB value is a result of 1 - accuracy. Furthermore, analyses were performed at genus level as well as at ZOTU level in order to detect associations that would not be displayed when grouping single sequences into large genus categories. All the remaining graphs were constructed with the R package “ggplot2” (version 3.4.0).

#### Data Availability

DNA sequencing data is available via the Sequence Read Archive (SRA) of the National Center for Biotechnology Information (NCBI): Accession number PRJEB61788.

## 3. Results

### 3.1. Overview

We analysed biofilm samples from 85 shower hoses originating from a single residential building using 16S and 18S amplicon sequencing for microbiome composition and ddPCR for *Legionella spp and L. pneumophila* relative abundance. These data were then analysed for potential associations between *Legionella* and members of the biofilm microbiome.

### 3.2. Microbial ecology of shower hoses biofilms

Shower hose biofilms fed with non-chlorinated drinking water consist of complex and diverse prokaryotic and eukaryotic communities (Figure 1). Amplicon sequencing revealed 1518 ZOTUs (16S, prokaryotic) and 1389 ZOTUs (18S, eukaryotic), respectively, with 111 prokaryotic and 78 eukaryotic taxa classified at genus level. These numbers represent a gross underestimation of the true diversity, since unclassified taxa at genus level accounted on average for 64.4±1.4% of the prokaryotic community and 85.3±1.2% of the eukaryotic community. Among the prokaryotic genera, *Meiothermus* spp. (3.6±0.6%), *H16* spp. (2.5±0.2%) and *Caulobacter* spp. (3.2±0.6%) were the most abundant, with the top ten organisms representing 19.6% of the identified community at the genus level. The eukaryotic community was composed of 10 phyla, of which *Amoebozoa* (7.8±0.9%)*, Excavata* (3.2±0.4%) and *Opisthokonta* (4.0±0.6%) were most abundant. Among the eukaryotic community, an average of 78±1.3% of the taxa were unclassified even at phylum level (in contrast, 0% of the prokaryotic community was unclassified taxa at phylum level). At the genus level, the three most abundant known eukaryotic taxa were *Hartmannella* (4.5±0.9%), *Limnofilidae_X* (1.6±0.3%) and *LKM74_lineage_X* (1.6±0.2%).

**Figure 1.**
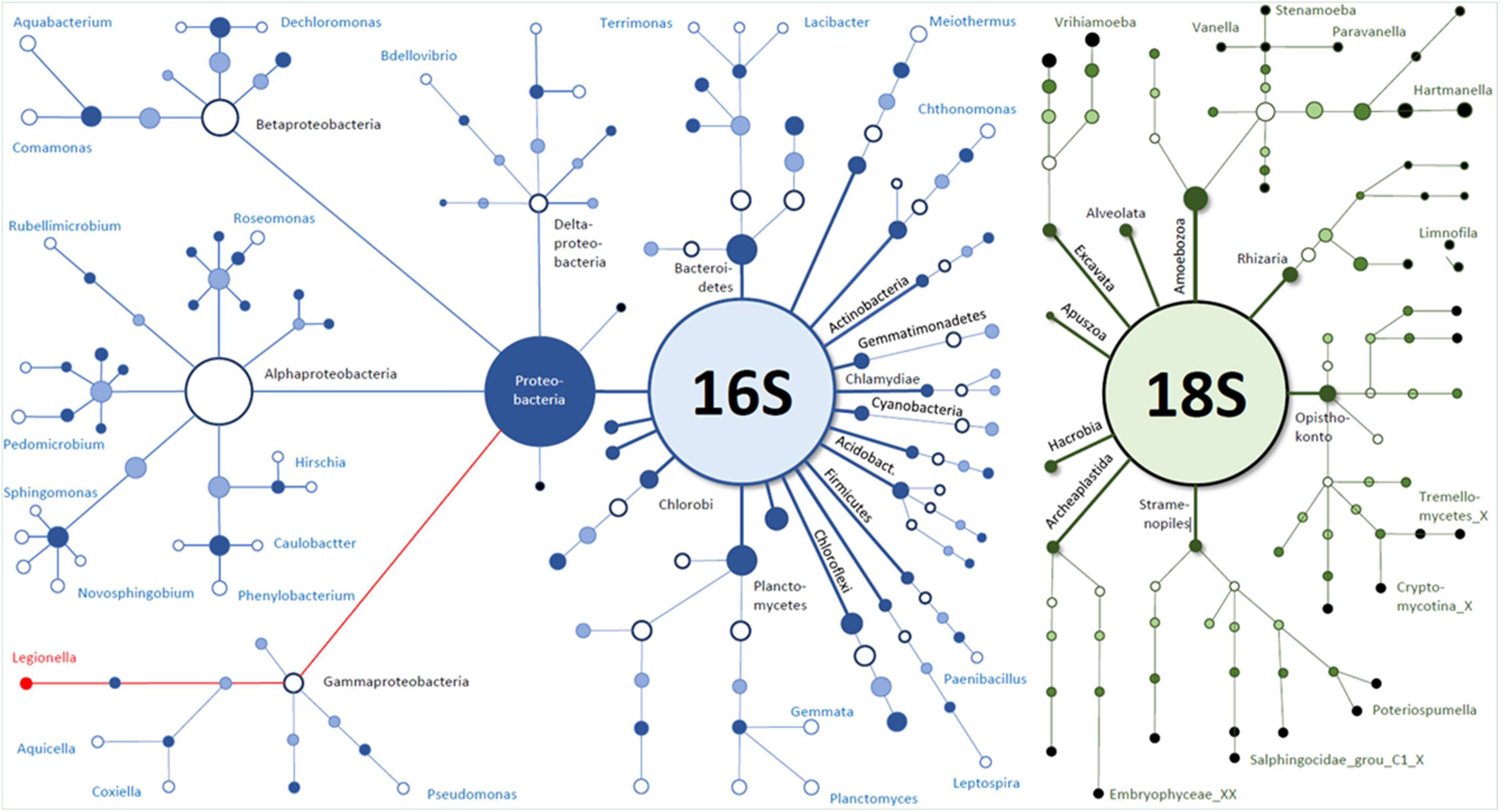
Overview of the average microbial composition of the 85 shower hose biofilms collected from a single building for this study. The figure shows average 16S and 18S data (n = 85) for ZOTUs above 0.01% relative abundance. Size of the nodes depicts the relative abundance of the corresponding taxa, while the connectors show the connection to the lower taxonomic level (from phylum to genus). Unclassified taxa at genus level account for 64.4±1.4% of the prokaryotic community and 85.3±1.2% of the eukaryotic community.

The distance between samples was measured using the Bray-Curtis index for both 16S and 18S data (Figure 2). In both cases, the samples did not show significant clustering with respect to the limited metadata available for this study (such as the location in the building where the sample campaign took place), indicating that the overall microbial composition of the samples was similar, with a few notable exceptions. However, relative abundance of individual taxa at genus level shows noticeable variations among samples (Supplementary Figure 1). In both 16S and 18S data, 12 samples (four shared; S001, S004, S008, S101) clustered separately from the bulk of the samples. This was likely due to the low richness of these samples. Alpha-diversity analysis for the 16S dataset shows high H values within samples, ranging between 3 and 5, with 11 samples having less than 3. The same analysis performed on the 18S dataset shows that most samples range between 3 and 4.8, while 12 samples have values below 3 (“H”, Supplementary Figure 2).

**Figure 2:**
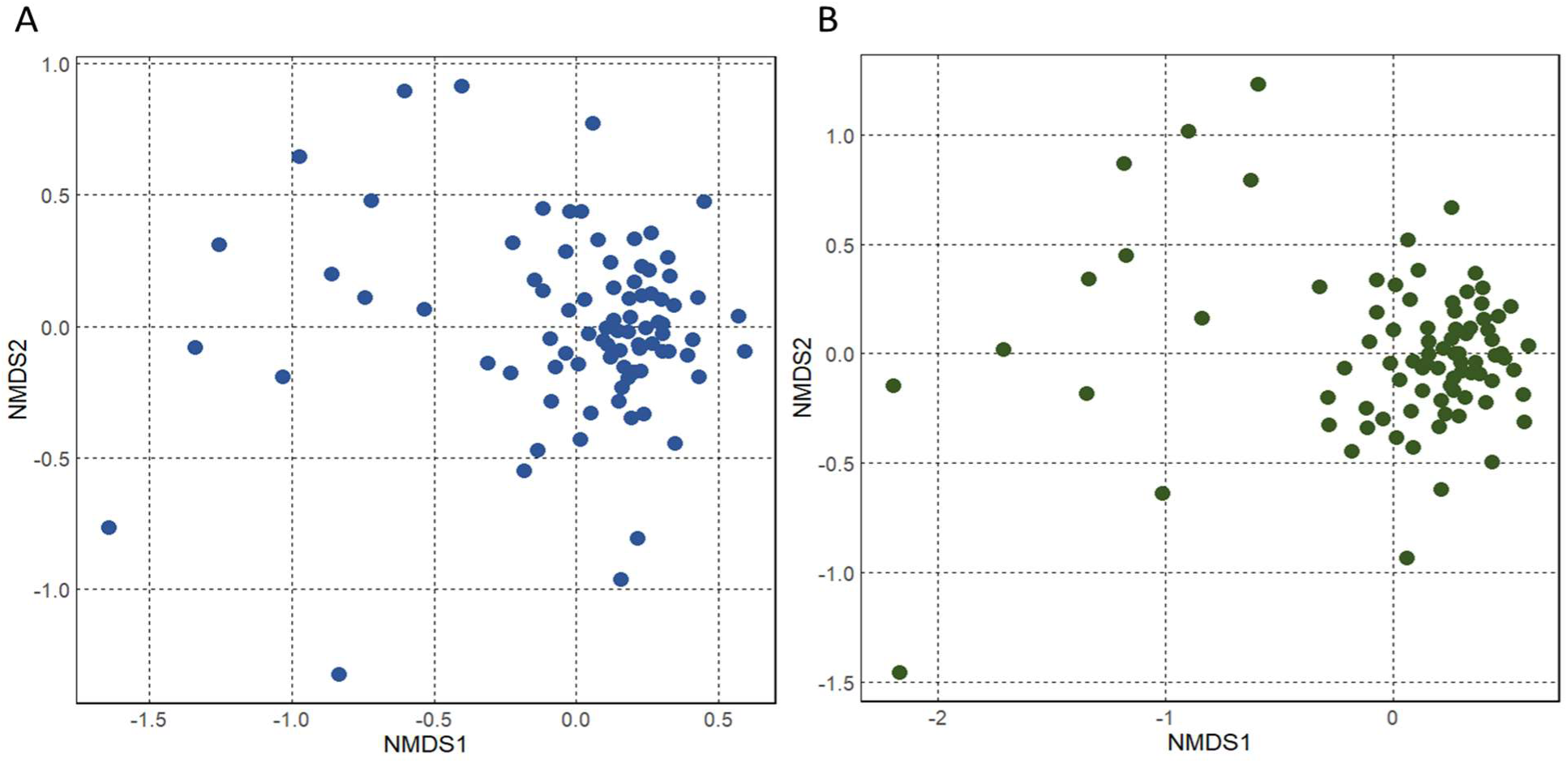
Ordination analysis of the (A) 16S and (B) 18S rRNA amplicon sequencing data from 85 shower hose biofilm samples, stress = 0.189 The graphs show distance among samples, calculated using the Bray-Curtis method and plotted on NMDS graphs.

### 3.3. Legionella spp. and L. pneumophila have different relative abundances across samples

The genus *Legionella* was a predominant taxon in many samples. With an average relative abundance of 0.27±0.02%, it ranked among the 30 most abundant genera in the community. Notably, 30 unique ZOTUs associated with the genus *Legionella* (the highest number of ZOTUs for a genus in this dataset) were detected and the relative abundance of these varied considerably across samples from the same building (Supplementary Figure 3).

We performed ddPCR for more specific quantification of *Legionella* spp and *L. pneumophila*, as well as for total 16S rRNA gene abundance (Figure 3). The relative abundance was observed across samples. *Legionella* spp. (relative abundance: 0.001% to 7.8%) was detected in all samples, while *L. pneumophila* (relative abundance: 0 to 0.46%) was detected in 34% of samples (Figure 3A). No clear correlation was observed between the relative abundance of *Legionella* spp. and *L. pneumophila*. Importantly, the relative abundance of *Legionella* spp. calculated from the 16S rRNA amplicon sequencing and calculated with ddPCR correlated poorly (R^2^=0.18; Figure 3B). Unless stated otherwise, the following analysis uses the targeted ddPCR data.

**Figure 3:**
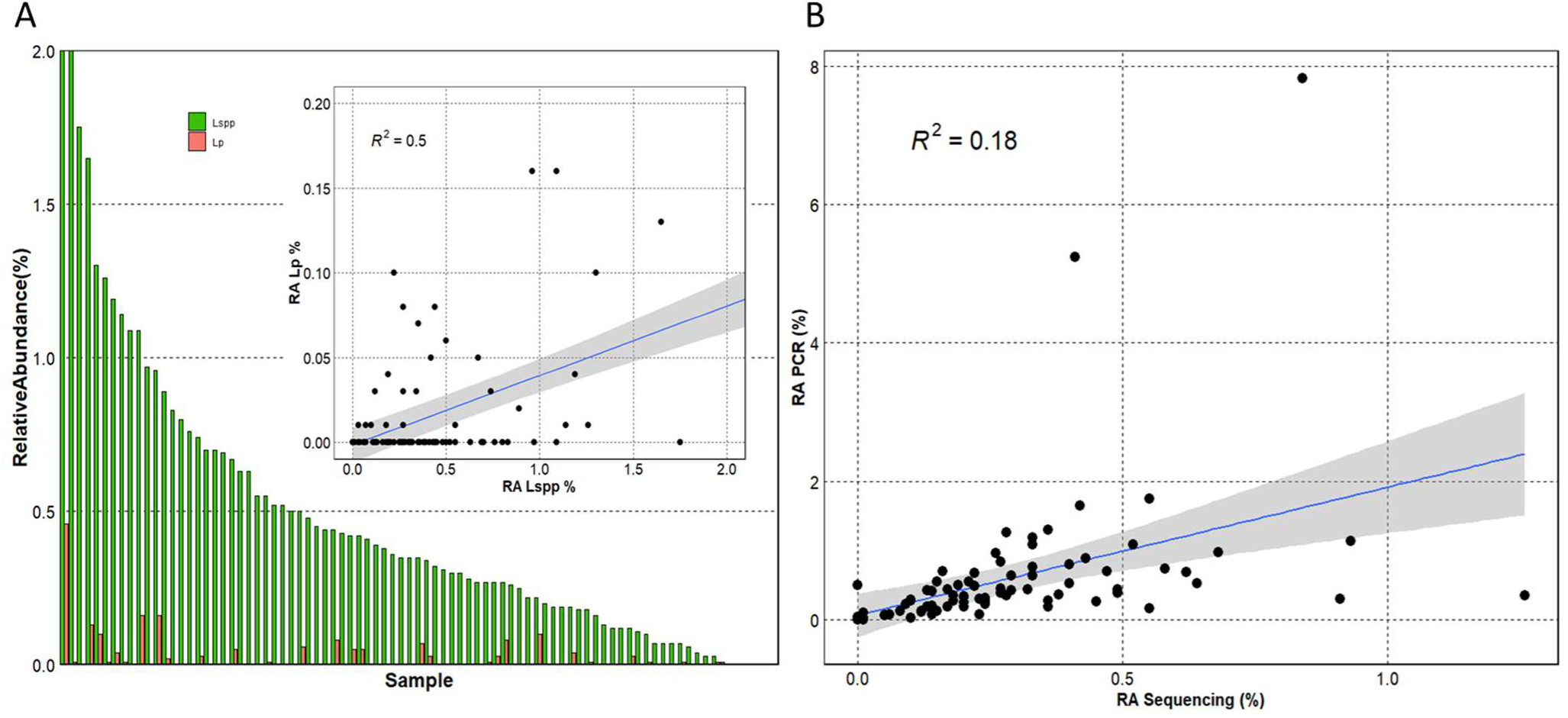
(A) Relative abundance of *Legionella* spp. (green bars) and *Legionella pneumophila* (orange bars) measured with ddPCR. Absolute quantification of *Legionella* spp. and *L. pneumophila* was measured with a duplex ddPCR assay, and concentrations were then normalized relative to the total 16S rRNA gene concentrations. The inlay figure displays the linear correlation (r = 0.5) between the relative abundance of *Legionella* spp. and *L. pneumophila* among samples (RA=Relative Abundance); (B) Linear correlation plot of the relative abundance of *Legionella* spp. calculated both with ddPCR and sequencing data.

### 3.4. Associations between *Legionella* spp. and members of the community

Random forest analysis identified combinations of prokaryotic and eukaryotic taxa that predicted the presence of high and low relative abundance of *Legionella* spp., but the choice of the model threshold affected the prediction (Figure 4, Supplementary Figures 4, 5). Using a median relative abundance threshold of 0.347% (equalling 42 samples with “high” relative abundance and 43 samples with “low” relative abundance), the model (at genus level) showed an OOB error of 0.275 for the 16S dataset (Figure 4A) and 0.43 for the 18S dataset (Figure 4E). At the ZOTU level, the model recorded an OOB error of 0.325 for the 16S (Figure 4B), and 0.376 for the 18S (Figure 4F) datasets. Main bacterial predictors of high relative abundance of *Legionella* spp. across models included *Chthonomonas*, *Planctomyces,* and *Sorangium*, with MDAs between 0.01 and 0.015 (Figure 4A), as well as ZOTU103 (g_*Reyranella*), ZOTU193 (g_*Legionella*), ZOTU11 (g_*Chthonomonas*) and ZOTU13 (g_*H16*) with MDAs between 0.003 and 0.004. Only two predictors of low *Legionella* spp. relative abundance, namely *Sphingopyxis* (MDA close to 0) and ZOTU 199 (MDA 0.0025; g_*Aquabacterium*) were detected. For the 18S dataset (Figure 4E), more predictors of low *Legionella* spp. abundance were observed (9 out of 15 predictors). The genera *Vanella* (linked to low abundance, MDA 0.008), *Vrihiamoeba* (linked to high abundance, MDA 0.008), and *Protacanthamoeba* (linked to high abundance, MDA 0.006) were the three main classified eukaryotic predictors of *Legionella* spp. relative abundance. ZOTU 58 (unclassified) predicted low *Legionella* abundance with an MDA of 0.025.

**Figure 4.**
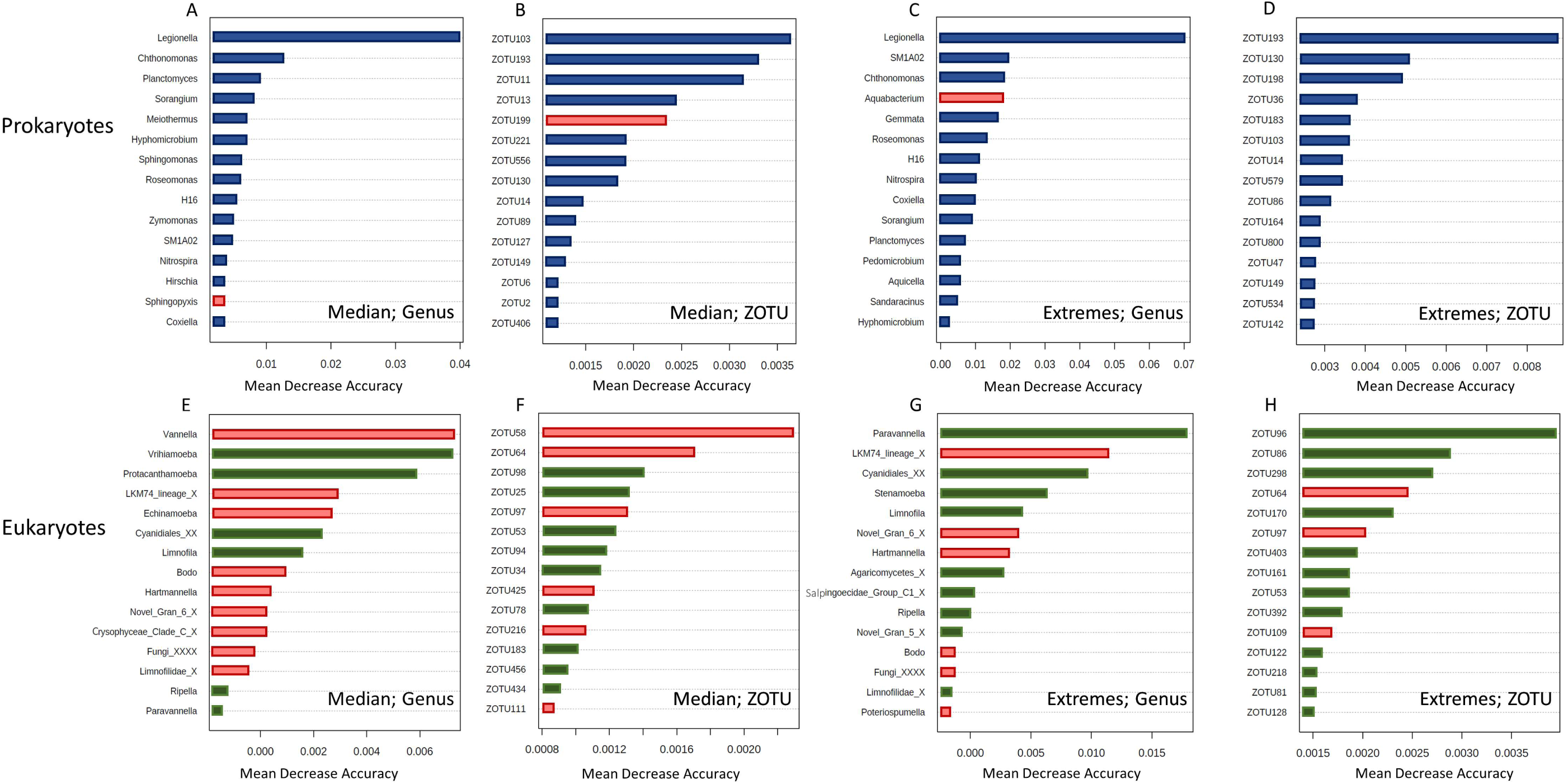
Random Forest analysis of the 16S and 18S datasets using the relative abundance of *Legionella* spp. as variable. Red represents predictors of low *Legionella* abundance while blue and green represent predictors associated with high abundance for prokaryotes and eukaryotes respectively. A-B: Main prokaryotes (genus level, A; ZOTUs level, B) predicting *Legionella* spp. abundance with a threshold of 0.3%; C-D: Main prokaryotes (genus level, C; ZOTU level, D) predicting *Legionella* spp. abundance using the samples at the extremes. E-F: Main eukaryotes (genus level, E; ZOTU, F) predicting *Legionella* spp. abundance using 0.3% as threshold; G-H: Main eukaryotes (genus level, G; ZOTU, H) predicting *Legionella* spp. abundance using the samples at the extremes.

However, selection of the criteria, such as the range of relative abundances to analyse for the model, is critical. When performing the analysis only using the samples at the extremes (15 samples with highest relative abundance; 15 samples with lowest relative abundance), the model returned an OOB error of 0.225 (16S) and 0.567 (18S) at the genus level and 0.2 (16S) and 0.5 (18S) at the ZOTU level (Figure 4A-D). For the prokaryotic community, *SM1A02* and *Chthonomonas,* consistently with the previous model, predict high *Legionella* abundance with MDAs of 0.0175, as did ZOTU193 (g_*Legionella*, MDA 0.01), ZOTU130 (c_*Acidobacter subgroup*, MDA 0.007) and ZOTU198 (g_*Coxiella*, MDA 0.005). *Aquabacterium* was the only trait that predicted low *Legionella* abundance (MDA 0.15.; Among the eukaryotic predictors, the genera *Paravannella* and *LKM74_lineage_X* consistently with the models described above, predict respectively high and low *Legionella* spp. abundance (MDA 0.0175; MDA 0.012). Three predictors of low *Legionella* spp. abundance are detected at the ZOTU level (ZOTU64; ZOTU97; ZOTU109, unclassified sequences). ZOTU96 (unclassified) was the stronger predictor of high *Legionella* spp. relative abundance, with an MDA of 0.004. We furthermore tested the models decreasing the thresholds between samples with high and low *Legionella* relative abundance to 0.2% (27 samples with low relative abundance; 58 samples with high relative abundance) and 0.1% (17 samples with low relative abundance; 68 samples with high relative abundance), which generated groups with uneven sample distribution and displayed different associations (Supplementary Figures S4-S5).

### 3.5. Associations between *L. pneumophila* and members of the biofilm microbiome

We also identified predictors of *L. pneumophila* relative abundance using Random Forest analysis. Using the median value (0.002%, Figure 3) as a threshold for samples with high and low relative abundance of *L. pneumophila*, the model performed with an OOB of 0.4 (16S) and 0.529 (18S) at genus level, and with OOB of 0.352 (16S) and 0.435 (18S) at ZOTU level. In contrast to the observations for *Legionella* spp. (above), the analysis showed more predictors of low abundance (eight at genus level; seven at ZOTU level). *Rhodobacter* (MDA of 0.014) and ZOTU57 (c_TK10, MDA 0.017) predicted low *L. pneumophila* relative abundance, while ZOTU52 (g_*Pedomicrobium*; MDA 0.030) was the most important predictor of high relative abundance. Only 10 features were detected with three negative associations, of which *LKM74_lineage_X* resulted as the most important (MDA 0.007). At the ZOTU level for the 18S dataset, the first three predictors are all associated with high *L. pneumophila* numbers (ZOTU342; ZOTU181; ZOTU168, unclassified sequences), while six ZOTUs predicted low *L. pneumophila* abundance, of which ZOTU230 (unclassified) had the highest importance.

We furthermore performed the analysis using only the extreme samples (25 samples highest abundance, 25 samples lowest abundance; Figure 3). Here the model performed with an OOB of 0.38 and 0.5 at the genus level, and 0.32 (16S) and 0.46 (18S) at the ZOTU level. For the 16S data, the results were similar to the previously described model, with more predictors of low *L. pneumophila* relative abundance (10). Interestingly, the model was not able to associate any predictors to the abundance of *L. pneumophila* for eukaryotes using these model parameters. At the ZOTU level, the prokaryotic ZOTU869 (g_*Legionella*) had the strongest importance value and an association with high *L. pneumophila* abundance. In general, more prokaryotic ZOTUs (11) predicted high *L. pneumophila* abundance in this model. For the eukaryotic sequences, the first three important ZOTUs predict low *L. pneumophila* abundance (ZOTU134; ZOTU48; ZOTU64, unclassified sequences).

### 3.6. *Legionella* ZOTUs correlations in a SparCC network analysis

SparCC analysis revealed several correlations between specific *Legionella* ZOTUs and other taxa, using only the 16S amplicon sequencing data. Figure 6 shows only significant correlation coefficients with a threshold of ±0.3, with a minimum of −0.36 and a maximum of 0.63. Since not all ZOTUs are assigned to the same level of taxonomy, the lowest level available was used to describe our observations. Of the 30 sequences associated to the *Legionella* genus, 18 ZOTUs did not display any significant correlation with other microbiome members above the threshold. The remaining 12 ZOTUs correlated in different ways with other sequences. For example, ZOTU193 was the most prevalent *Legionella* sequence in the dataset (Supplementary Figure 3) and was also the one displaying the most correlations (67), the majority of which were positive. Only four negative correlations have been detected, with ZOTU104 (g_*Ferribacterium*), ZOTU765 (f_*Obscuribacterales*), ZOTU156 (g_*Lacibacter*), and ZOTU1309 (g_*Rhodoplanes*). The strongest positive correlation (0.63 correlation coefficient) is with ZOTU33 (g_*H16)*.

The analysis also showed that *Legionella* ZOTUs associated in different ways with the same taxa. For example, while the *Legionella* ZOTU193 positively correlated with ZOTU135 (o_*Obscuribacterales*) and ZOTU196 (f_*Rhodobiaceae*) while *Legionella* ZOTU1258 displayed negative correlations with these. The same phenomenon was observed for ZOTU55 (g_*SM1A02*, positively correlates with *Legionella* ZOTU193, negatively correlates with *Legionella* ZOTU1483) and for ZOTUs 323 and 37, (f_*Planctomycetaceae*; g_*Pedomicrobium*; positively correlate with *Legionella* ZOTU193 and negatively correlate with *Legionella* ZOTU1379). ZOTU996 mostly displayed negative correlations (nine, ranging from −0.3 to −3.6): the only positive correlation (coefficient of 0.322) was with ZOTU817 (g_*Bdellovibrio*), while ZOTU78 (g_*Nitrospira*), ZOTU347 (g_*H16*) were the ones with the highest negative correlation coefficients. In only two cases, two *Legionella* ZOTUs correlated with each other, in both cases positively: ZOTU735 and ZOTU640 correlated with each other (correlation coefficient 0.3), as well as ZOTU462 and ZOTU241 (correlation coefficient 0.31). The genus *Chthonomonas was always* associated with high *Legionella* spp. abundance and positively correlated with the *Legionella* ZOTU801 (correlation coefficient 0.3)

## 4. Discussion

Community analysis of 85 shower hose samples revealed a highly diverse prokaryotic and eukaryotic composition (Figure 1) that was relatively consistent among samples (Figure 2). However, *Legionella* spp. and *L. pneumophila* relative abundance varied considerably among samples (Figure 3). Various Random Forest analyses demonstrated that several prokaryotic and eukaryotic taxa were associated with either high or low relative abundance of *Legionella* spp. and respectively *L. pneumophila* (Figures 4-5). Moreover, SparCC correlation analysis showed how individual *Legionella* ZOTUs correlated with specific microbiome members (Figure 6).

**Figure 5.**
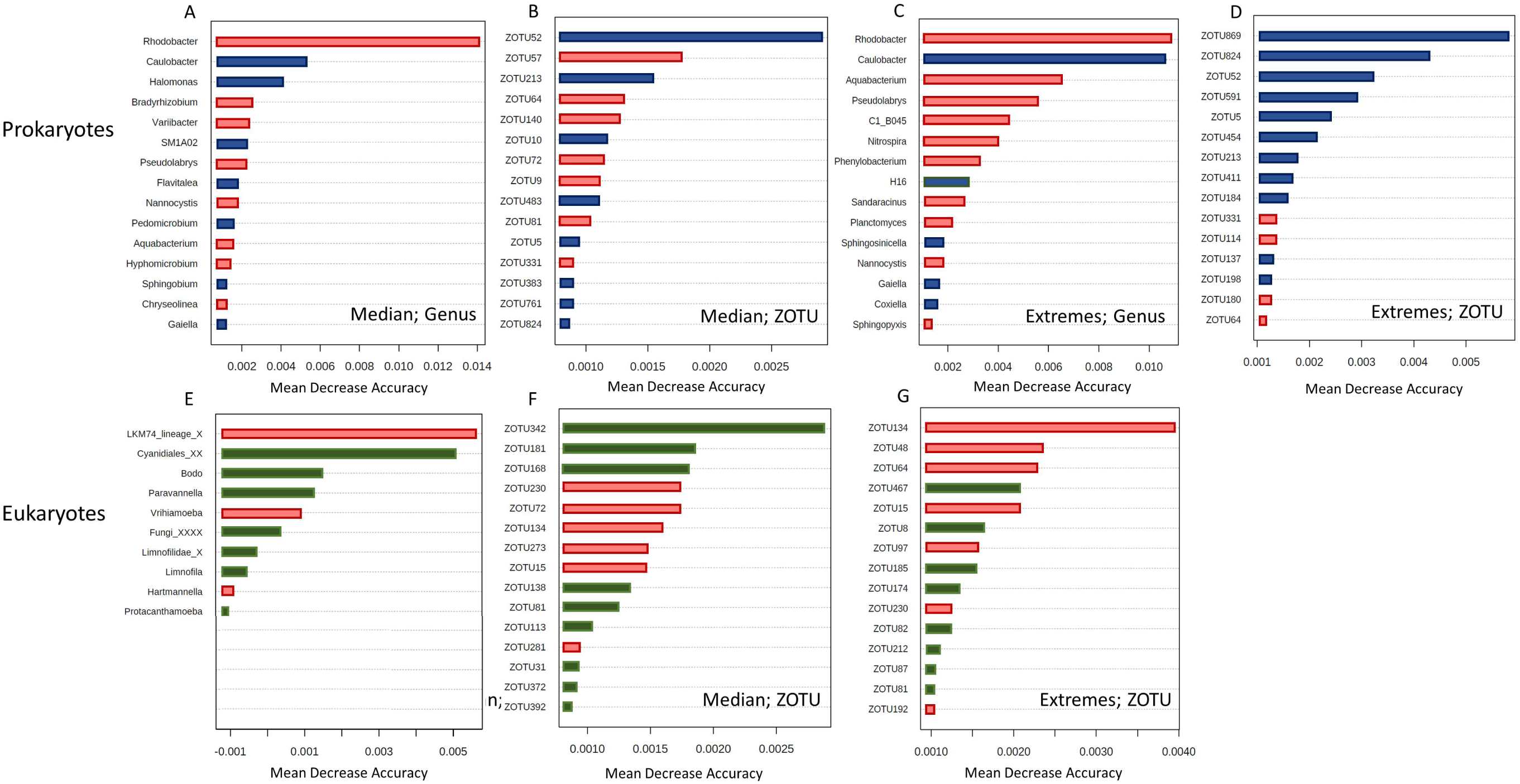
Random Forest analysis of 16S and 18S datasets using the relative abundance of *Legionella pneumophila* as variable. Red represents predictors of low Legionella abundance while blue and green represent predictors associated with high abundance for prokaryotes and eukaryotes respectively. A-B: Main prokaryotes (genus level, A; ZOTUs level, B) predicting *L. pneumophila* abundance with a threshold of 0.02% relative abundance; C-D: Main prokaryotes (genus level, C; ZOTU level, D) predicting *L. pneumophila* abundance using the samples at the extremes. E-F: Main eukaryotes (genus level, E; ZOTU, F) predicting *L. pneumophila* abundance using 0.02% relative abundance as threshold; G-H: Main eukaryotes (ZOTU, G) predicting *L. pneumophila* abundance using the samples at the extremes.

### 4.1. Microbial ecology in the final meters of water distribution

Shower hoses biofilms are inhabited by diverse prokaryotic and eukaryotic communities. Three main factors promote microbial growth here: 1) temperature, which is often warmer in the last meters; 2) flexible materials that leach organic carbon; 3) usage patterns that often create extensive stagnation periods (Proctor *et al*., 2016). This makes shower hoses ideal environmental niches where biofilms form and microorganisms can proliferate. Differences in the above-mentioned factors also means that there is considerable diversity between different shower biofilm microbiomes (Proctor *et al*., 2016, Gebert *et al*., 2018). To mitigate the uncertainty caused by such differences, we collected all hoses on purpose from a single building, which shared the same incoming water, had a similar age, and were made from the same material. The shower hose biofilms showed general similarity in terms of common diversity indices, with dominant taxa similar to those identified from previous studies of non-chlorinated drinking water systems *(Liu et al., 2014, Proctor et al., 2018, Neu et al., 2019)*. The average relative abundance of the microorganisms is highly variable among biofilm samples (Figure S1), as well as the relative abundance of *Legionella* spp. and *L. pneumophila,* and *Legionella*-associated ZOTUs (Figure 3, Supplementary Figure 3). While the reasons behind this variability are not clear, the data provided interesting information: 1) *Legionella* spp. relative abundance reaches very high values in shower hoses biofilms (7.8% in our samples); 2) the higher relative abundance of *Legionella* spp. compared to *L. pneumophila*, as well as the 30 ZOTUs classified as *Legionella*, suggest multiple *Legionella* species inhabiting individual shower hose biofilms. Despite the lack of available studies, the presence of multiple *Legionella* species with different abundances has been detected in different water systems, although poorly described in shower hoses to our knowledge (Lesnik *et al*., 2016, Logan-Jackson & Rose, 2021, Salinas *et al*., 2021, Gleason *et al*., 2023).

Many studies cover prokaryotic drinking water biofilms, but information on eukaryotes in these systems is still severely limited. Studies focussing on eukaryotes detected amongst others the fungal sub-phylum *LKM11* and the amoeba clade *LKM74* in biofilm samples of different DWDSs, together with other organisms like *Streptophyta*, Ciliates and Algae, *Opisthokonta, Stramenopiles and Alveolata* (Inkinen *et al*., 2019, Soler *et al*., 2021). We also detected these taxa, but, importantly, as well as several taxa that are known carriers of opportunistic pathogens, such as *Vannella* spp., *Hartmannella* spp., *Acanthamoeba* spp. and *Echinamoeba* spp. (Inkinen *et al*., 2019). However, the relevant eukaryotic community is likely under-represented with current 18S amplicon sequencing tools. For example, 18S primers target all eukaryotes, which increases the chances of detection of undesired species/contaminations. In response, specific databases have been developed for certain fractions of eukaryotes of interest to the exclusion of others (Guillou *et al*., 2013). A further major drawback is that the taxonomic identification of eukaryotes is still limited and this calls for improvements in all aspects of eukaryotes ecology research, from targeted sample collection to dedicated analysis pipelines (Gabrielli *et al*., 2023).

All this information together highlight how shower hoses biofilms are environments full of diversity, where interactions happen from species to kingdom level (Sadiq *et al*., 2022). As shown, *Legionella* spp. are members of these environments, and therefore subjected to different ecological interactions.

### 4.3. Some prokaryotic taxa are associated with *Legionella* abundance

Microbes in biofilms exist in close proximity, thus creating opportunities to establish positive and/or negative interactions with each other. Random Forest analysis showed a prevalence of positive associations between the resident microbiomes and *Legionella* spp. / *L. pneumophila* (Fig. 4 A-D; Fig. 5 A-D). Positive bacterial interactions include promoting the growth of other bacteria by increasing general nutrient availability, creating new niches or directly exchanging important nutrients (e.g., cross feeding of amino acids) (D’Souza *et al*., 2018, Kehe *et al*., 2021). In the context of *Legionella* growth, such beneficial interactions can play a decisive role: *Legionella* are auxotrophs for at least seven amino acids (Tesh & Miller, 1981, Chien *et al*., 2004).

In contrast, negative associations between prokaryotes and *Legionella* can include direct (interference) or indirect (exploitative) competition (Granato *et al*., 2019, Cavallaro *et al*., 2022). Our results suggest a negative association between *Legionella* spp. and only two genera, namely *Aquabacterium* and *Sphingopyxis* (Figure 4). It is not possible to exactly establish the nature of these interactions using only amplicon sequencing data. While these two taxa are not known to be producers of secondary metabolites with antimicrobial activity, they are characterized by a highly versatile metabolism, which provides advantages with respect to surviving to unfavourable environmental conditions (Sharma *et al*., 2021). This can in theory give them an advantage in terms of competition for resources in low-nutrient environments compared to other microorganisms, especially the ones with specific and high nutrient requirements, like *Legionella*.

A few previous studies have explored associations between *Legionella* and the resident microbiome in different systems. In two studies focusing on cooling towers, Paranjape and colleagues found bacteria associated with the presence or absence of *Legionella* spp. and *L. pneumophila* (Paranjape *et al*., 2020, Paranjape *et al*., 2020). *Pseudomonas* was found as the main genus correlating with the absence of *Legionella*, as already reported elsewhere (Leoni *et al*., 2001, Proctor *et al*., 2018). Interestingly, our study did not show a strong negative association between *Pseudomonas* and *Legionella*. In addition, Paranjape and co-workers reported *Sphingopyxis* as a positive correlating genus, in contradiction with our results (Figure 4). A recent investigation on plumbing systems across four cities of Europe revealed associations between culturable *Legionella* and the microbial communities detected in the water samples (Scaturro *et al*., 2022). While reinforcing the negative association between *Legionella* and *Pseudomonas*, the authors also described a negative association with the genera *Staphylococcus* and, consistently with our analysis, *Sphingopyxis*.

Associations between bacterial communities and culturable *L. pneumophila* in hot and cold water were also investigated in a study conducted on high rise buildings in New York City (Ma *et al*., 2020). In that study, associations were described at higher taxonomic levels and negatively linked *L. pneumophila* and members of the classes *Bacteroidia* and *Solibacterales*, while positive associations were detected with *Chloracidobacteria*, *Gemm-1,* and *Actinobacteria*, among others. The fact that disagreement exists between the various studies depends on different factors. For example, previous studies analysed exclusively the water phase, in contrast with our study that focused on the biofilm phase. Moreover, the taxonomic classification at the genus (sometimes class) level does not allow for specific identification of the organisms involved, and different species within the same genus might exhibit different associations with the same organism. We explored the latter point by looking at the correlations that specific *Legionella* ZOTUs exhibit in a SparCC analysis (Figure 6; discussed below). The real ecological meaning of these associations is not clear, as correlation can obviously not be linked automatically to causation. Many microbial ecology studies aim to find associations between two species that reflect their ecology in the environment (Carr *et al*., 2019). However, interactions between species often occur in high-order combinations, where an interaction between two species is modulated by a third one (Bairey *et al*., 2016). Therefore, associations detected by statistical analysis may not necessarily describe direct interactions between microorganisms. It is also possible that two organisms are correlating with a common environmental variable rather than each other. Nevertheless, these observational studies create good opportunities to further study bacterial interactions using laboratory approaches. For example, Paranjape et al. (Paranjape *et al*., 2020) isolated a strain of *Brevundimonas* and confirmed the positive effect of the strain towards the growth of *L. pneumophila* previously observed through statistical analysis.

**Figure 6.**
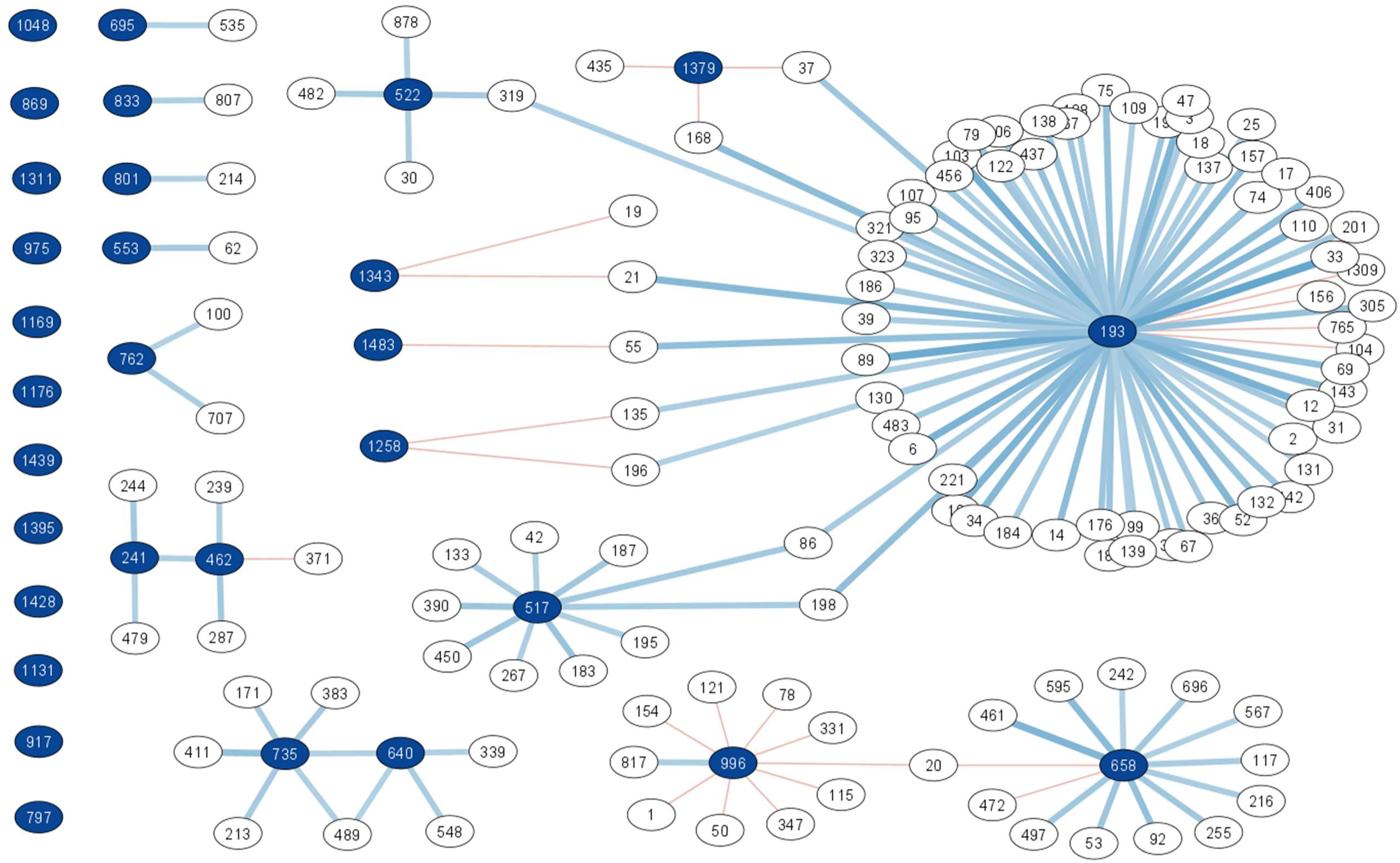
SparCC analysis constructed as a network using the 16S amplicon sequencing dataset, subsetted for the ZOTUs associated with *Legionella* taxonomy. To help interpretation, the nodes corresponding to the *Legionella* ZOTUs are coloured in dark blue, while the edges are in light blue indicating positive correlations and orange for negative correlation coefficients. In the top right are all the *Legionella* ZOTUs that do not show any significant correlation (above 0.3 correlation coefficient).

### 4.4. Some eukaryotic taxa are also associated with *Legionella* abundance

Random forest analysis performed on the eukaryotic data set revealed more negative associations with *Legionella* abundance compared to the prokaryotic data. Unravelling the meaning of these associations is challenging, as this is likely linked to the intracellular lifestyle of *Legionella*. Interactions between protists and *Legionella* can, in fact, have very different outcomes. For example, *Legionella* are taken up by their host as a potential food source, but escape the host’s degradation mechanisms and start the replicative phase by establishing in the *Legionella*-containing-vacuole (LCV) (Boamah *et al*., 2017). When this happens, the protist hosts are often killed (thus resulting in increased *Legionella* abundance and decreased host abundance). Alternatively, some protists effectively graze on *Legionella* as a food source and are resistant to their escape mechanisms, resulting in decreased *Legionella* abundance and increased host abundance (Dey *et al*., 2009, Amaro *et al*., 2015, Boamah *et al*., 2017, Mondino *et al*., 2020). Paranjape and co-workers interpreted a positive association between *Legionella* and *Oligohymenophorea* as an indication of host-prey relationship in which *Legionella* is able to replicate (potentially without causing the death of the host) (Paranjape *et al*., 2020). To add to the complexity, it has been reported that the temperature and the abundance of *Legionella* can change the fate of host-prey relationships, which can be either digested, packaged into vesicles and secreted in pellets, or survive intracellularly with no replication (Boamah *et al*., 2017).

Different *Legionella* species can also be associated with different outcomes with the same host (Boamah *et al*., 2017). Our dataset identified several associations with protists proposed as *Legionella* hosts. *Vannella* spp., *Echinamoeba* spp. and *Hartmannella* spp. (now *Vermamoeba*) predicted the low abundance of *Legionella* in the biofilm samples. While *Vannella* has been previously described as able to kill *Legionella* (Rowbotham, 1986), *Vermamoeba* is often referred to as a permissive host for *Legionella*, although cases of intracellular survival have been reported (Buse & Ashbolt, 2011). To our knowledge, no information on how the interaction *Echinamoeba*-*Legionella* takes place is available. Additional negative associations include *Novel-Gran-6*, which are members of the phylum *Cercozoa*, described as particularly prone to digest *Legionella (Boamah et al., 2017).* The genera *Vrihiamoeba* and *Protacanthamoeba* predicted, on the other hand, high *Legionella* abundance. No clear interaction between these taxa and *Legionella* has to our knowledge been described, but both are known to prey on bacteria (Delafont *et al*., 2014, Ma *et al*., 2016). Interestingly, *Vrihiamoeba* is a predictor of low *L. pneumophila* abundance, indicating that the same interaction could produce a different outcome depending on the specific *Legionella* species involved.

### 4.5. Associations are often interactions that are happening at small taxonomic scale

A limitation of community analysis performed through 16S amplicon sequencing is that genus-based generalization of the interactions is usually made, while species dynamics is not considered. In response, we explored the diversity and associations of *Legionella* ZOTUs (Figure 6; Supplementary Figure 3). Previous studies discussed the possibility to identify species of *Legionella* using only a partial sequence of the 16S rRNA gene (Wilson *et al*., 2007, Ma *et al*., 2020). If correct, this offers an opportunity to study ecological dynamics of the genus *Legionella* at species level in environmental samples.

While this evidence combined suggest that different species of *Legionella* inhabit building plumbing biofilms, the Random Forest analysis used either the entire genus or specifically *L. pneumophila* datasets as external variables for the predictions. To challenge this further, we focused on the correlations of the individual *Legionella* ZOTUs, which suggested that the 30 sequences display markedly different associations in the complex ecological network represented by these biofilms. A first observation of this network, consistently with the Random Forest data, highlights a prevalence of positive associations between *Legionella* and the resident microbiome. In the specific case of this study, different *Legionella* sequences have dissimilar correlation patterns, and correlations between individual *Legionella* ZOTUs are rare. This suggests the possibility that *Legionella* species live within different and defined ecological niches in biofilms, and that they interact/associate with different microorganisms (Rottjers & Faust, 2018). This is, from an ecological point of view, an important aspect as more than 25 species within the genus *Legionella* are considered pathogenic and we believe that studying the ecology of non-pneumophila species is as important.

### 4.6. Critical considerations of the methods used to detect associations

The associations identified in this study display similarities, but also differences with other studies in the field (discussed above). This is influenced by a number of factors, which are not always clearly highlighted in similar studies:

1. *Legionella quantification*: We used both ddPCR and 16S amplicon sequencing, and results from the two methods did not correlate particularly well (Figure 3B). Moreover, both these methods would amplify and detect non-viable *Legionella*, of which the importance in ecological-association studies is questionable. Previous studies with a similar focus used cultivation-based quantification methods (Ma *et al*., 2020, Scaturro *et al*., 2022), which also has known bias towards *L. pneumophila* and is limited by only analysing cells that grow on agar plates (Toplitsch *et al*., 2021). Clearly, the method of *Legionella* quantification would considerably affect the outcome of any correlation/association analysis.
2. *Tools used to determine associations*: Changing the threshold on the relative abundance of *Legionella* (high *Legionella* vs low *Legionella*) affected the outcome of the Random Forest models (Supplementary Figure 5). This is partially due to the OOB errors of the models, which are especially higher for the prediction of low *Legionella* abundance. From a biological point of view, this is an important consideration: choosing the median relative abundance value as the threshold is statistically stronger as all the samples are part of the Random Forest analysis. Focusing only on the samples at the extremes would, however, in theory define more relevant predictors of the presence/absence of *Legionella*. In addition, different statistical tools (i.e., SparCC and Random Forest analysis) predictably did not produce the same result but have complementary value. The Random Forest analysis identifies general predictors of the abundance of *Legionella* spp. and *L. pneumophila* in a given environment; SparCC analysis provides us with evidence on the separate ecological niches that different *Legionella* species occupy. In the former, the cause for high OOB values indicates the model is less accurate than desired while the latter highlights that associations of microorganisms ideally need to be resolved at a smaller taxonomic and spatial scale. Other studies have used linear regression or LefSe analysis to link *Legionella* to other microorganisms (Paranjape *et al*., 2020, Paranjape *et al*., 2020). Our methodology choice can be justified by the data collected in our study: targeted ddPCR allows us in fact to use the relative abundance of *Legionella* as an external variable that can be predicted by the amplicon sequencing results (validated moreover by the fact that *Legionella* is displayed as a predictor of high *Legionella* abundance). While not advocating for certain methods over others, it is obvious that the choice of method would affect the result. An ideal situation would be standardizing methods and protocols across different studies, and reporting as much metadata as possible, as to gain generalizable and comparable insights into *Legionella* ecology and microbial associations in diverse environments.
3. *Biofilm vs water phase:* In our study, the biofilm phase was specifically collected and analysed, compared to other investigations where the water phase was analysed (Ma *et al*., 2020, Paranjape *et al*., 2020, Scaturro *et al*., 2022). While noting the challenges in biofilm sampling, we do believe that this choice is critical for the determination of associations between microorganisms. The microbial composition of biofilm and water is different and not necessarily reflective of each other (Chan *et al*., 2019), and therefore associations detected in the two phases are likely different. Also, *Legionella* are clearly recognised as biofilm-associated bacteria (Declerck, 2010), hence, associations within this spatial niche should be most relevant. Finally, the bacterial composition of established biofilms bacterial composition is more stable in time, while water is a more dynamic system (Inkinen *et al*., 2014).

### 4.7. Function vs taxonomy

A strong limitation of conventional amplicon sequencing is that only taxonomic information is obtained after processing, and it is often not possible to resolve the dataset at species level. This does not allow a good overview of the species dynamics in given samples, while making it also nearly impossible to look at specific functions shaping the ecology of the environments (Lajoie & Kembel, 2019). Shotgun metagenomics may be one alternative approach (Eisen, 2007, Perez-Cobas *et al*., 2020). The advantages of such a method compared to amplicon sequencing are a finer taxonomic resolution (it is possible in some cases to reconstruct entire genomes) and, more importantly, the detection of genes with their abundances) associated with specific functions. This would benefit investigations on *Legionella* ecology and interactions with other organisms as well. Our study identified several organisms correlating with/predicting the abundance of *Legionella* and we explored potentially explanations for these interactions. Using shotgun metagenomics in these studies would, inform on the specific functions associated with the abundance of *Legionella*, which would in turn help creating a more detailed overview on what characterizes an environment as favourable or undesirable for *Legionella* proliferation, being moreover able to link the genes to the organisms. We previously proposed potential probiotics approaches against *Legionella* in engineered aquatic ecosystems (Cavallaro *et al*., 2022), and reviewed studies that identified antagonistic behaviours of certain microorganisms. The use of shotgun metagenomics in observational studies can be a useful resource in this sense, as the identification of specific functions that make the environment undesirable for *Legionella* can be searched for in specific microorganisms to use in such an approach.

## 5. Conclusions

- Non-chlorinated shower hose biofilms from the same building plumbing system are inhabited by diverse prokaryotic and eukaryotic communities, including widely varying relative abundances of *Legionella*.
- Eukaryotic communities are important for a complete understanding of the ecology of *Legionella* in plumbing systems, but a considerable expansion of 18S databases is crucial for the identification of key organisms and associations.
- A combination of prokaryotic and eukaryotic organisms statistically predicted the abundance of *Legionella* spp. (and *L. pneumophila*), but the direct ecological meaning of specific associations is ambiguous. The low predicting quality of the models, however, needs to be considered when putting the associations in a biological context.
- Analysing the associations looking at single amplicon sequence variants (rather than genus) gives an overview of potential interactions at species level, and a deeper ecological understanding of *Legionella* species dynamics.
- Moving towards function-based approaches (i.e. metagenomics of drinking water biofilms) will provide a better understanding of the mechanisms that affect *Legionella* abundance. This will in turn allow the use of specific known functions as predictors of *Legionella* abundance.

## Supporting information

Report 16S data processing

Report 18S data processing

## Acknowledgements

This research was funded by the Federal Food Safety and Veterinary Office (FSVO), in partnership with the Federal Offices of Public Health (FOPH) and Energy (SFOE) in Switzerland, through the project LeCo (Legionella Control in Buildings; Aramis nr.:4.20.01) and Eawag discretionary funding.

Data produced and analyzed in this paper were generated in collaboration with the Genetic Diversity Centre (GDC), ETH Zurich.

