## Supplementary Information for "*Legionella* relative abundance in shower hose biofilms is associated with specific microbiome members"

Alessio Cavallaro^1,2^, William J. Rhoads^1^, Emile Sylvestre^1^, Thierry Marti^1,2^, Jean-Claude Walser^2^, Frederik Hammes^1^*

^1^ Department of Environmental Microbiology, Eawag, Swiss Federal Institute of Aquatic Science and Technology, 8600 Dübendorf, Switzerland

^2^ Department of Environmental Systems Science, Institute of Biogeochemistry and Pollutant Dynamics, ETH Zurich, 8092 Zurich, Switzerland

* Corresponding author:

Name:	Frederik Hammes

**Table of Contents**

**Supplementary Figure 1.** Average Relative Abundance of the 30 top taxa classified at genus level	**3**

**Supplementary Figure 2.** Alpha diversity of the 16S and 18S datasets

**4**

**Supplementary Figure 3.** Relative Abundance of the individual *Legionella* ZOTUs across samples	**5**

**Supplementary Figure 4.** Thresholds used for Random Forest analysis on *Legionella* spp. and *L. pneumophila* relative abundance	**6**

**Supplementary Figure 5.** Random Forest analysis on *Legionella* spp. relative abundance using additional thresholds	**7**

**Supplementary Table 1.** Primers, probes and ddPCR reagents and conditions for a duplex assay to detect *Legionella* spp. and *Legionella pneumophila*	**8**

**Supplementary Table 2.** Primers, probes and ddPCR reagents and conditions for the detection of the total 16S genes	**9**

**Supplementary Table 3.** Primers, and PCR reagents and conditions for the amplicon sequencing library preparation	**10**

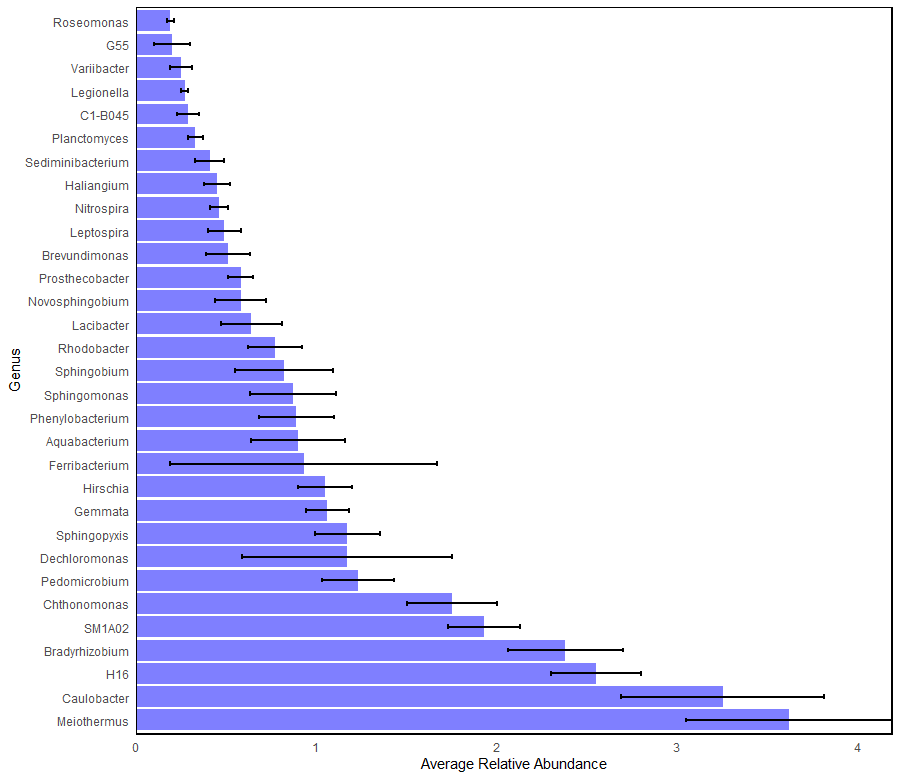

**Supplementary Figure 1. Average Relative Abundance of the 30 top taxa classified at genus level**

The figure shows the average relative abundance across samples of the 30 most abundant taxa classified at genus level. The relative abundance is shown in increasing order from the less abundant to the most abundant, according to the absolute value. A standard error has been included in the plot to show the variations across samples. The relative abundance has been calculated using the ZOTU count table produced with the 16S rRNA gene amplicon sequencing.

A

B

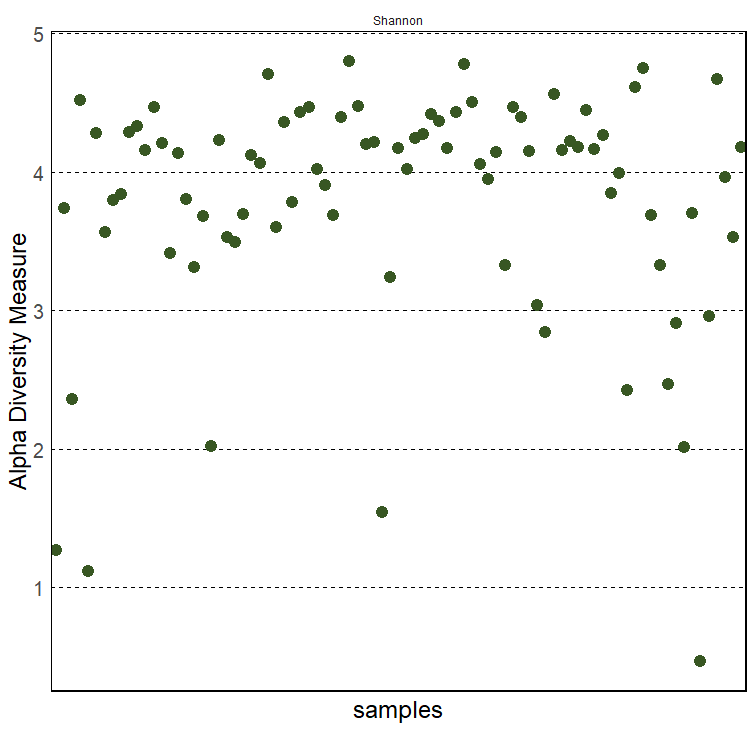

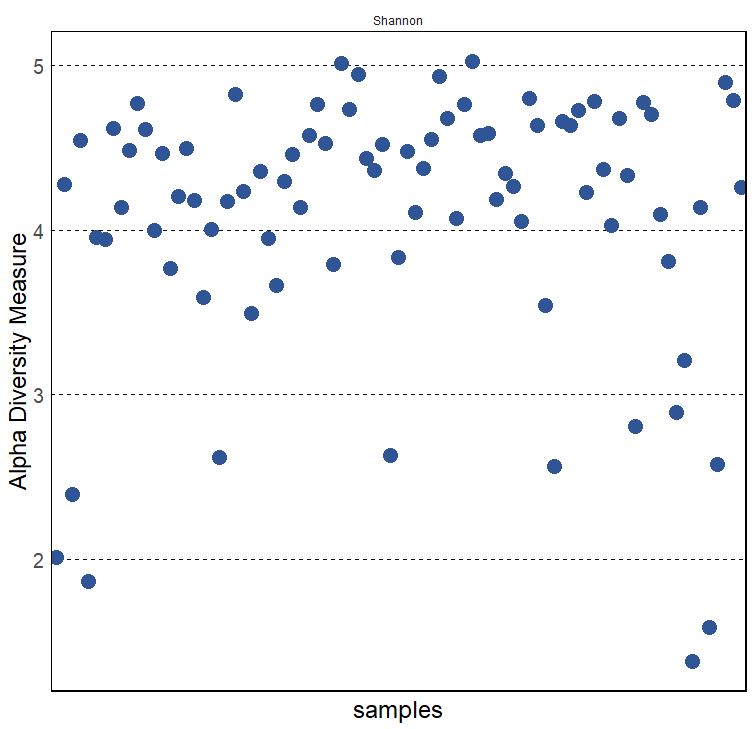

**Supplementary Figure 2. Alpha diversity of the 16S and 18S datasets**

The alpha diversity for the 16S and 18S rRNA gene amplicon sequencing datasets has been calculated using the Shannon index, which takes into account both richness and evenness. The Shannon index (H) is indicated in the y-axis, and the intra-sample diversity is considered higher as the H index increases.

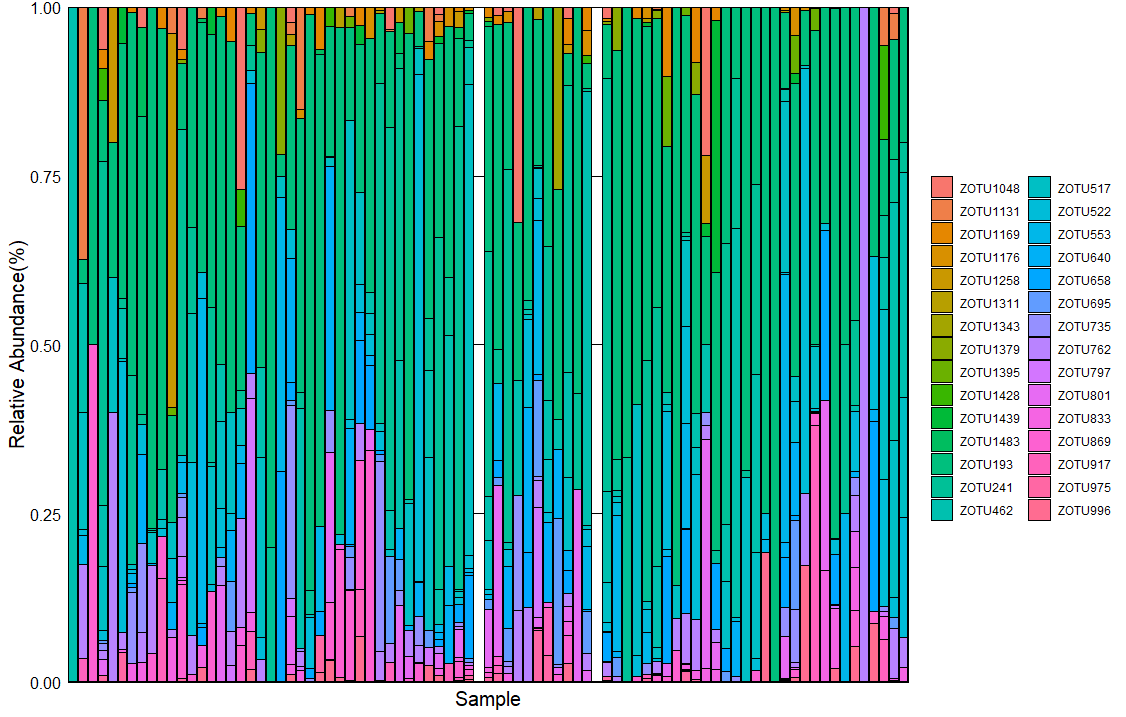

**Supplementary Figure 3. Relative Abundance of the individual *Legionella* ZOTUs across samples**

This stacked bar plot displays the distribution of the 30 ZOTUs assigned to the genus *Legionella* across multiple samples, with each bar representing a single sample. The bar height reflects the total abundance of the genus within each sample, while the segments within the bar indicate the relative abundance of individual ZOTUs. This visualization provides insight into the diversity and relative importance of different ZOTUs within the *Legionella* genus across samples.

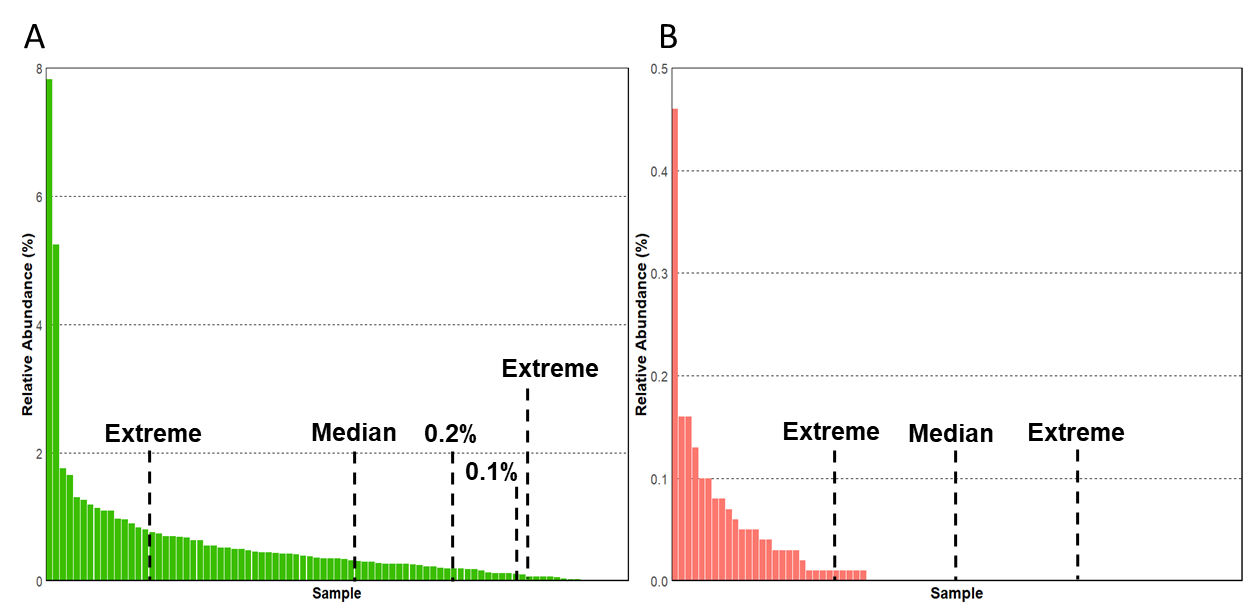

**Supplementary Figure 4. Thresholds used for Random Forest analysis on *Legionella* spp. and *L. pneumophila* relative abundance**

The figure shows how the different thresholds used to determine the variables in the Random Forest analysis were applied to the relative abundance of *Legionella* spp. and *L. pneumophila*. A) The selection of the thresholds for *Legionella* spp. relative abundance. The median value (0.347%) creates two groups of 43 and 42 samples, while only 15 samples per side were choses as extremes based on their relative abundance. Using a threshold of 0.2% results in an uneven sample distribution, with 27 low-*Legionella* samples and 58 high-*Legionella* samples. Further decreasing the threshold to 0.1% creates two groups of 17 and 68 samples. B) Selection of the thresholds for L.pneumophila. The median value (0.002%) creates two groups of 39 and 46 samples (three samples have a relative abundance of 0.002% and have been all included in the same group). 25 samples at the extremes were chosen for this analysis, in order to accommodate all the samples with no *L. pneumophila* in one group.

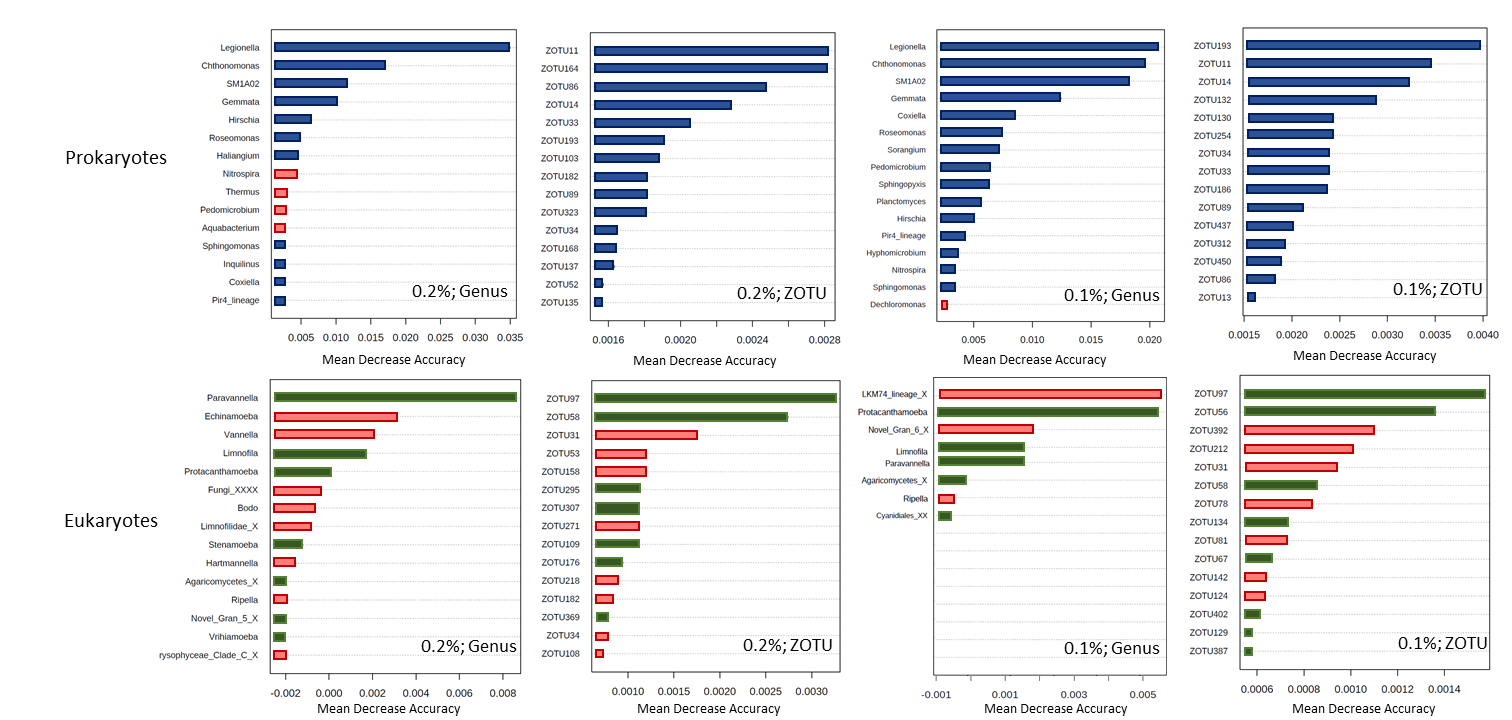

**Supplementary Figure 5. Random Forest analysis on *Legionella* spp. relative abundance using additional thresholds**

Random Forest analysis of 16S and 18S datasets using the relative abundance of *Legionella* spp*.* as variable and additional thresholds (0.2% and 0.1% Legionella spp. relative abundance). A-B: Main prokaryotes (genus level, A; ZOTUs level, B) predicting *Legionella* spp. abundance with a threshold of 0.2% relative abundance; C-D: Main prokaryotes (genus level, C; ZOTU level, D) predicting *Legionella spp.* abundance with a threshold of 0.1% relative abundance. E-F: Main eukaryotes (genus level, E; ZOTU, F) predicting *Legionella* spp*.* abundance using 0.2% relative abundance as threshold; G-H: Main eukaryotes (ZOTU, G) predicting *Legionella* spp*.* abundance using a threshold of 0.1% relative abundance.

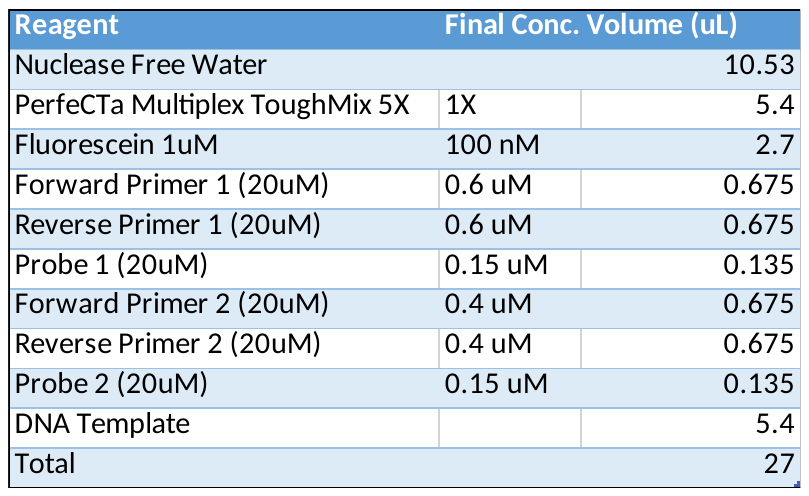

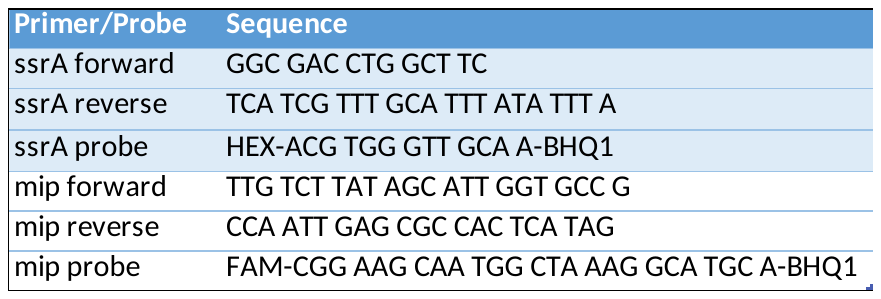

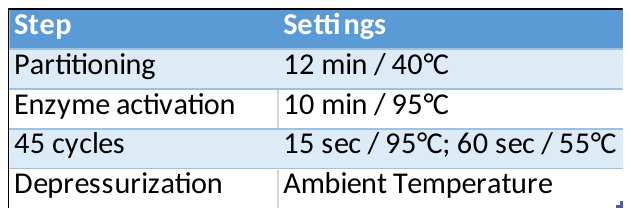

**Supplementary Table 1. Primers, probes and ddPCR reagents and conditions for a duplex assay to detect *Legionella* spp. and *Legionella pneumophila***

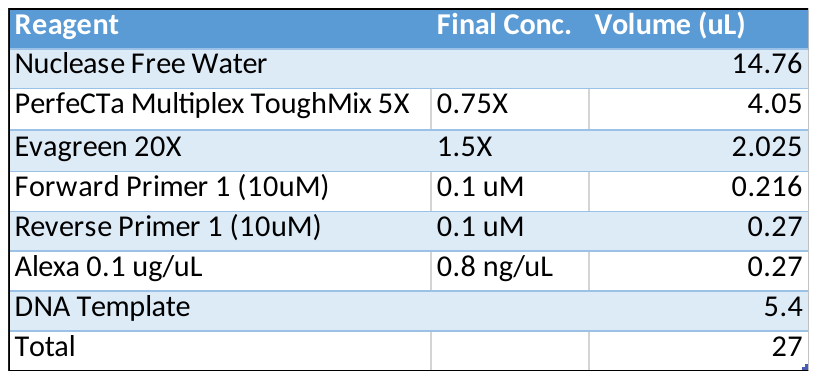

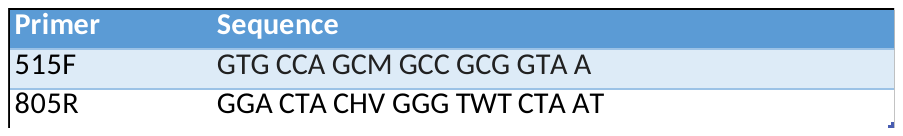

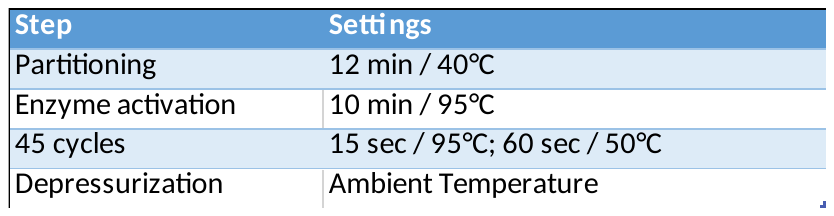

**Supplementary Table 2. Primers, probes and ddPCR reagents and conditions for the detection of the total 16S genes**

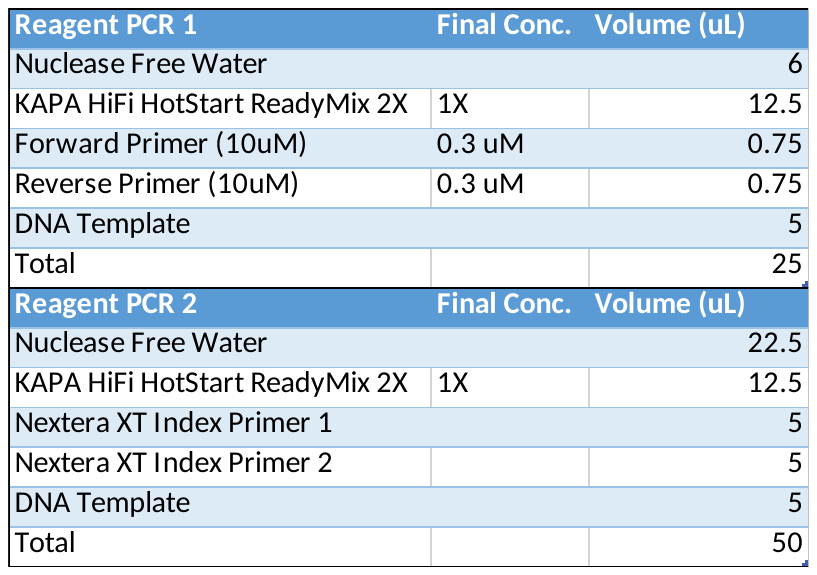

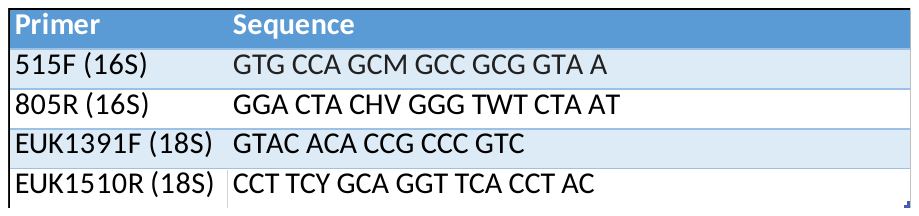

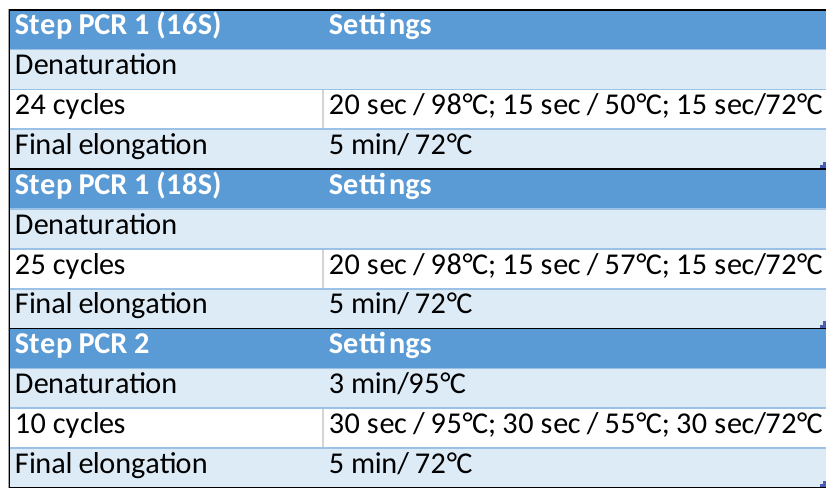

**Supplementary Table 3. Primers, PCR reagents and conditions for the amplicon sequencing library preparation**
